# Clustered Rapid Induction of Apoptosis Limits ZIKV and DENV-2 Proliferation in the Midguts of *Aedes aegypti*

**DOI:** 10.1101/2020.10.15.341370

**Authors:** Jasmine B. Ayers, Heather G. Coatsworth, Seokyoung Kang, Rhoel R. Dinglasan, Lei Zhou

## Abstract

Inter-host transmission of pathogenic arboviruses such as dengue virus (DENV) and Zika virus (ZIKV) requires systemic infection of the mosquito vector. Successful systemic infection requires initial viral entry and proliferation in the midgut cells of the mosquito followed by dissemination to secondary tissues and eventual entry into salivary glands^1^. Lack of arbovirus proliferation in midgut cells has been observed in several *Aedes aegypti* strains^2^, but the midgut antiviral responses underlying this phenomenon are not yet fully understood. We report here that there is a rapid induction of apoptosis (RIA) in the *Aedes aegypti* midgut epithelium within 2 hours of infection with DENV-2 or ZIKV in both *in vivo* blood-feeding and *ex vivo* midgut infection models. Inhibition of RIA led to increased virus proliferation in the midgut, implicating RIA as an innate immune mechanism mediating midgut infection in this mosquito vector.

---

ZIKV and DENV serotypes 1-4 (DENV-1-4) are arthropod-borne viruses (arboviruses) of the genus *Flavivirus* that cause acute febrile illness in humans. Upon reinfection with a different dengue viral serotype, antibody dependent enhancement can result in severe and potentially fatal manifestations such as hemorrhagic fever and dengue shock syndrome^3^. There are an estimated 100 million symptomatic dengue infections annually^4^. Although acute infection with ZIKV is typically asymptomatic or mild, it has been found to cause neurological sequelae such as Guillain-Barre syndrome in adults and neurological birth defects such as microcephaly in infants born to infected mothers^5,6^.

DENV and ZIKV are both primarily transmitted between humans by *Aedes aegypti*. The primary site of viral infection is the mosquito midgut epithelium, and the midgut is the first physical defense barrier against virus establishment. The proliferation of arboviruses in the midgut following consumption of an infected blood meal is a prerequisite for subsequent systemic infection of the vector and eventual transmission of the virus. Interestingly, not all *Ae. aegypti* have an equivalent ability to contract and transmit DENV and ZIKV^7^. Failure to establish viral infection in the midgut after a flavivirus infected blood meal has been observed in both field-derived lab *Ae. aegypti* strains such as Cali-MIB and in wild-collected individuals^2,8–10^. The mechanisms by which mosquitoes resist these pathogens are of great epidemiological interest, both for predicting the ability of local mosquito populations to transmit these viruses and for engineering mosquito strains that could resist infection.

Apoptosis has been posited as an important innate immune response against viral infection in both insects and mammals^11–13^. One line of evidence is that viral genes with anti-apoptotic function are crucial for infectivity of insects by several families of virus^14,15^. ZIKV in particular has recently been shown to encode subgenomic RNAs implicated in inhibiting apoptosis and increasing infectivity in mosquitoes^16^. In addition, expression of pro-apoptotic regulatory genes by genetically engineered Sindbis virus demonstrated that inducing apoptosis can limit viral infection of *Aedes aegypti17*. However, depending on the virus/vector pairing studied, apoptosis observed following arbovirus infection of mosquitoes correlates in some cases with refractoriness^16–21^ and in other cases with susceptibility^22–24^. Apoptosis in insects is in some cases a mechanism that facilitates viral release-in Rift Valley Fever virus infection of *Anopheles stephensi*, apoptosis appears to be necessary for viral escape into the salivary gland lumen^23^; the baculovirus *Autographa californica* M nucleopolyhedrovirus requires caspase activation to pass through the midgut basal lamina^25^, and *Spodoptera frugiperda* ascovirus encodes its own executioner caspase, which is expressed during late stages of the replication cycle^26^. The seemingly contradictory roles of apoptosis in insect virus infection suggest a balancing point between infection control and pathological tissue damage. The kinetics of the apoptotic response in relation to the viral life cycle appears to be an important factor in that balance. Previous work on apoptosis following viral infection in the adult mosquito midgut has focused on 24+ hours post infection^16–24^. However, experiments conducted in *Drosophila* indicate that a rapid induction of apoptosis (RIA), which occurs within 4 hours following injection of flock house virus (FHV) is responsible for limiting systemic infection^27^. The present study explores whether this RIA response and its limiting effect occurs in vector mosquitoes infected with medically relevant *Flaviviruses*.

Using adult female Orlando (ORL) strain *Aedes aegypti*, we assayed for signs of apoptosis following exposure to a naive bloodmeal or a bloodmeal containing either DENV-2 NGC (DENV serotype 2 New Guinea C strain) or ZIKV-PR (lab-adapted Puerto Rico ZIKV strain PRVABC59). At 2 hours after blood feeding began, the midguts were dissected and immediately fixed **(Fig. 1a)**. Using TUNEL (Terminal deoxynucleotidyl transferase dUTP nick end labeling) to detect the DNA fragmentation, we noted only a few apoptotic cells in the midgut epithelium of mosquitoes fed with a naive bloodmeal (**Fig. 1b)**. In contrast, there was a 29.1-fold increase (p = 3.74e-06) in mean TUNEL-positive cells in the midgut of mosquitoes that consumed a bloodmeal containing ZIKV (**Fig. 1c**), and a 19.6-fold increase in DENV-2 fed mosquitoes (p = 1.18e-06) (**Fig. 1d**). The few TUNEL-positive cells in naive blood-fed midguts were discrete and surrounded by cells that were negative for TUNEL (**Fig. 1b**). In contrast, in DENV-2 or ZIKV fed midguts, TUNEL-positive cells often appeared in clusters of 10-50 cells (**Fig. 1c, 1d**). The position of the TUNEL-positive cells in the midgut following virus feeding appears to be random and not restricted to particular regions of the midgut. An increase in TUNEL-positive cells in *in vivo* ORL midguts can be detected as soon as 1 hour following a ZIKV bloodmeal, and this elevation persists to 4 hours post-infection (hpi) **(Extended Data Figure 1**). At 8 hpi and 24 hpi, both naive blood fed mosquitoes and virus fed mosquitoes had an equivalent, high number of TUNEL-positive cells (mean of 465 TUNEL-positive cells/gut; **Extended Data Figure 1**), which is consistent with previous work showing a high level of midgut turnover in hematophagous mosquitoes during blood meal digestion^28^. We termed the rapid appearance of TUNEL-positive cells before 4 hpi as a rapid induction of apoptosis (RIA) to distinguish the phenomenon from the much larger, nonspecific induction of apoptosis observed at later time points.

**Fig. 1:**
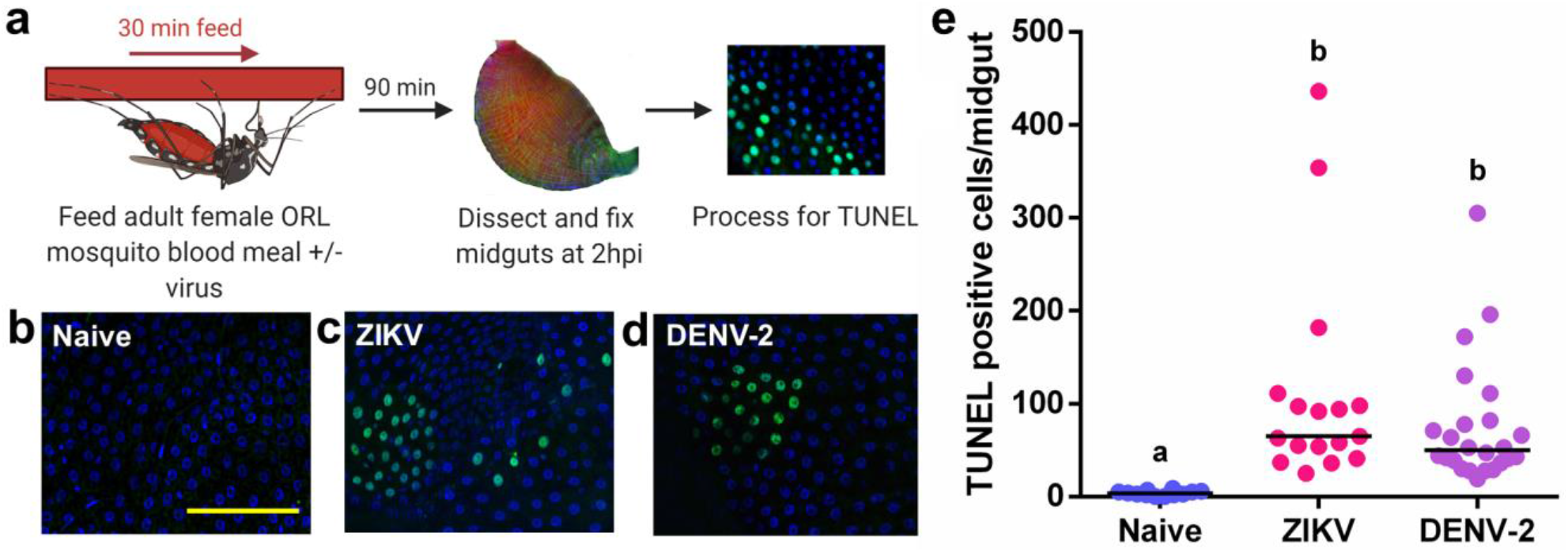
Exposure to a bloodmeal containing DENV-2 or ZIKV induces DNA fragmentation in the midgut epithelium of adult female *Aedes aegypti* (ORL) mosquitoes at 2hpi. **a,** Workflow diagram of *in vivo* infection. **b-d,** Representative images of DNA fragmentation visualized via TUNEL with DAPI counterstain in midguts from adult female ORL mosquitoes at 2 hours post **(b)** naive blood meal **(c)** ZIKV infected blood meal or **(d)** or DENV-2 infected blood meal. Scale bar = 100 μm. **e,** Quantification of TUNEL positive cells per midgut at 2 hours post *in vivo* infection (*n* (naive) = 12; *n* (ZIKV) = 17; *n* (DENV-2) = 24). The horizontal line indicates the median. Treatments without a common letter were found to be statistically significant (α = 0.05) as calculated by Kruskal-Wallis test with Mann-Whitney post hoc comparison (Kruskal-Wallis chi-squared = 32.017, df = 2, p-value = 1.116e-07).

To orthogonally verify that exposure to ZIKV or DENV-2 induces RIA in the midgut, we developed an *ex vivo* infection model, whereby midguts were dissected from adult female ORL mosquitoes that had not been blood fed and were incubated at room temperature in either media alone or media containing virus (10^6^ PFU/mL) **(Fig. 2a)**. At 2 hours following exposure to either ZIKV or DENV-2, there was a 9.8-fold increase in mean TUNEL-positive cells per midgut over the no virus control in ZIKV treated midguts (p = 3.59e-05) and a 9.1-fold increase in DENV-2 treated midguts (p = 0.000433) (**Fig. 2b-e**). The patterning of these TUNEL-positive cells was similar to what was seen in the *in vivo* infection model, as we frequently observed clusters of 10-50 cells with no obvious anterior to posterior preference in localization. In addition to DNA fragmentation, there was an extensive activation of caspases detected via a pan-caspase *in situ* activation reporter in the midgut 2 hours after DENV-2 or ZIKV exposure (**Fig. 2f-h**).

**Fig. 2:**
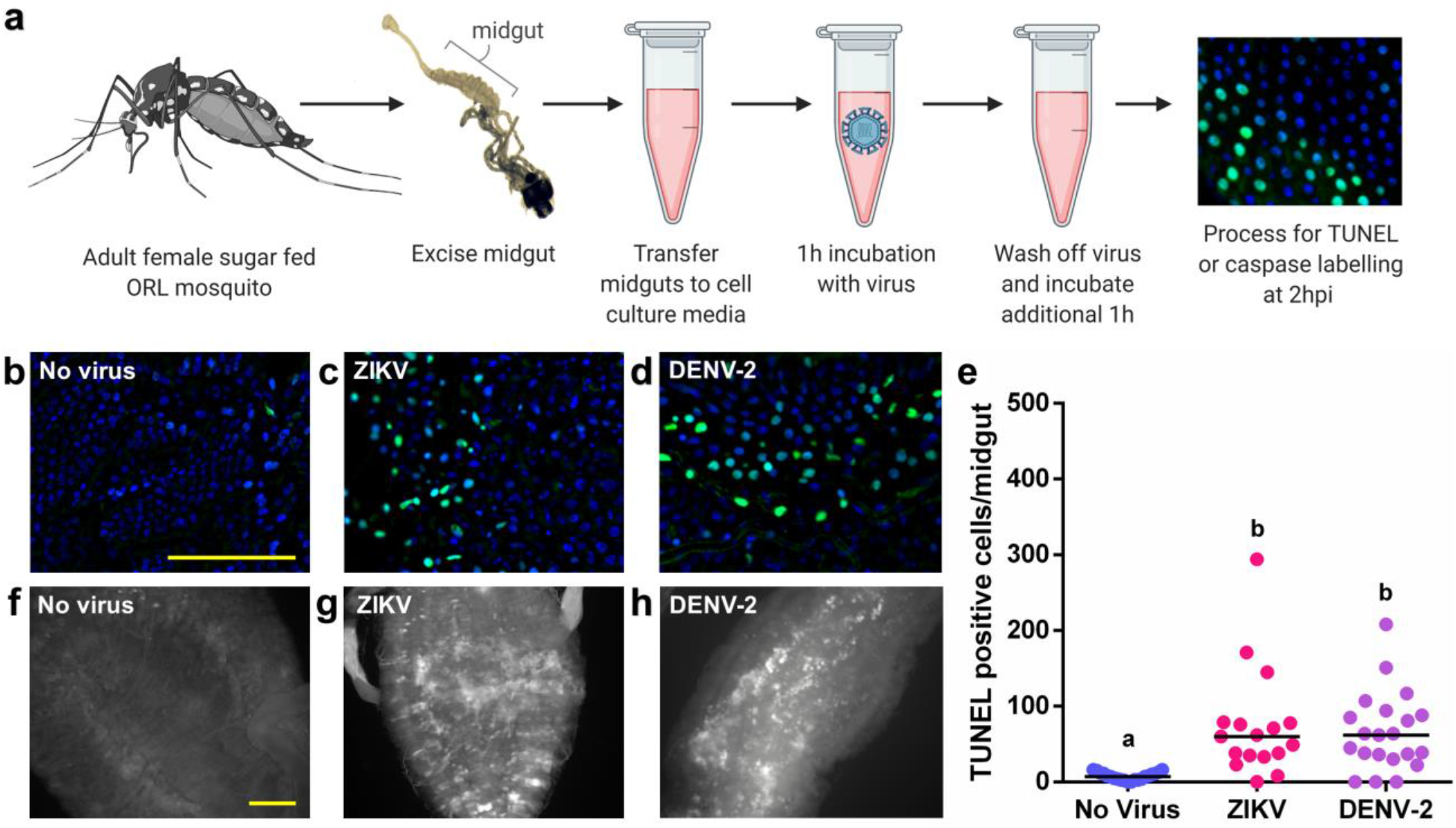
Exposure to a bloodmeal containing DENV-2 or ZIKV induces DNA fragmentation in the epithelium of *ex vivo* adult female *Aedes aegypti* (ORL) midguts at 2hpi. **a,** Workflow diagram of *ex vivo* midgut infection. **b-d,** Representative images of DNA fragmentation visualized via TUNEL at 2 hours after **(b)** *ex vivo* mock infection, **(c)** ZIKV infection or **(d)** DENV-2 infection. Scale bar = 100 μm. **e,** Quantification of TUNEL positive cells per midgut at 2 hours post *ex vivo* infection. (*n* (naive) = 21; *n* (ZIKV) = 17; *n* (DENV-2) = 18). The horizontal line indicates the median. Treatments without a common letter were found to be statistically significant (α = 0.05) as calculated by Kruskal-Wallis test with Mann-Whitney post hoc comparison (Kruskal-Wallis chi-squared = 22.285, df = 2, p-value = 1.448e-05). **f-h,** Pan-caspase activation at 2 hours after **(f)** mock infection **(g)** ZIKV infection or **(h)** DENV-2 infection. Scale bar = 100 μm.

To test whether the RIA response correlated with a refractory phenotype in strains with differential virus susceptibility, we chose the MOYO-Refractory (MOYO-R) and MOYO-Susceptible (MOYO-S) strains of *Ae. aegypti,* which were established with selective inbreeding from the original MOYO-In-Dry strain for their respective resistance and susceptibility to *Plasmodium gallinaceum* (avian malaria parasite)^29^; they were later found to be refractory (19.54% infection rate) and susceptible (56.60% infection rate), respectively, to DENV-2 (Jamaica 1409 strain) infection^30^. When both MOYO-R and MOYO-S mosquitoes were fed with a DENV-2-infected blood meal, there was a 1.8-fold higher mean number of TUNEL-positive epithelial cells in MOYO-R mosquitoes as compared to MOYO-S mosquitoes (p = 0.00212) (**Fig. 3**).

**Fig. 3:**
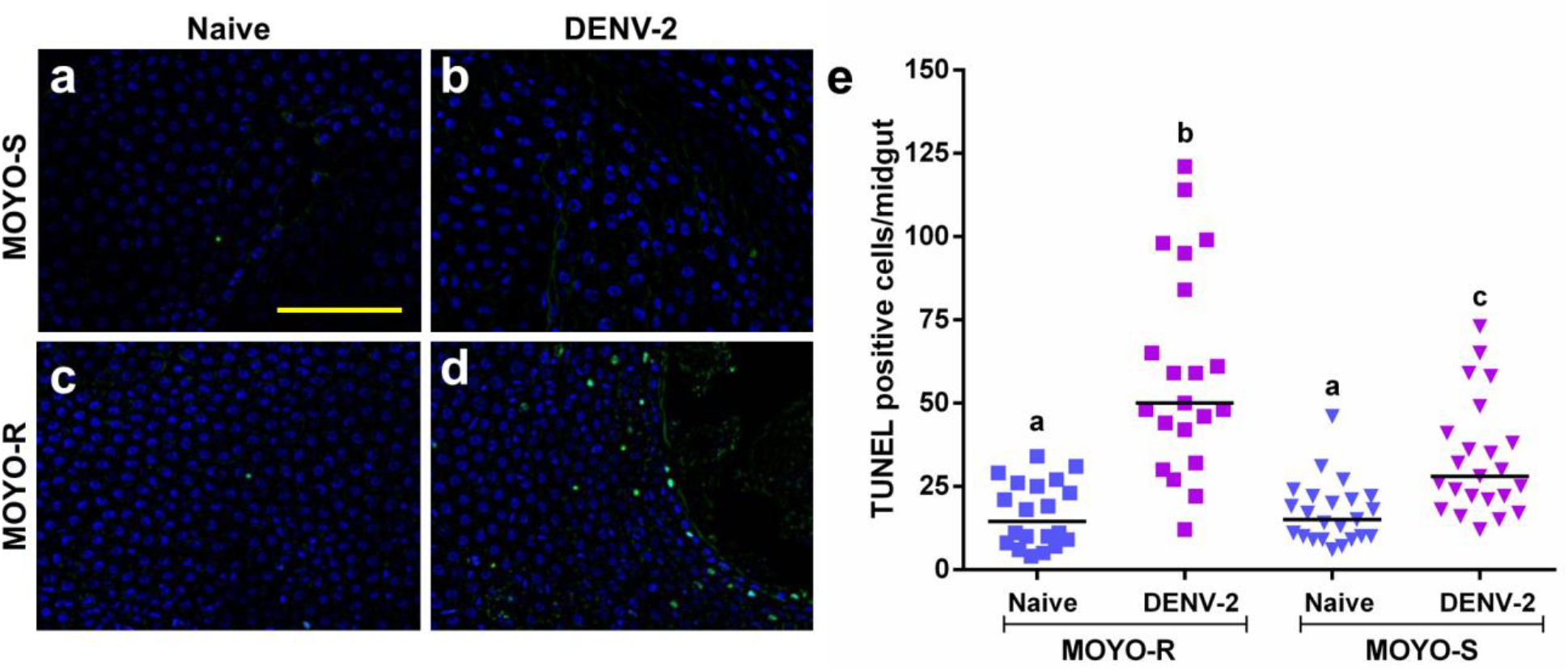
*Aedes aegypti* MOYO-R strain mosquitoes that are partially resistant to DENV-2 have higher rates of TUNEL-positive cells in the midgut than DENV-2 susceptible MOYO-S strain mosquitoes at 2 hours post ingestion of a DENV-2 infected blood meal. **a-d,** Representative images of TUNEL with DAPI counterstain in midguts from adult female mosquitoes at 2 hours post blood meal. **(a)** MOYO-S mosquitoes fed naive blood, **(b)** MOYO-S mosquitoes fed DENV-2 infected blood meal, **(c)** MOYO-R mosquitoes fed naïve blood or **(d)** MOYO-R mosquitoes fed DENV-2 infected blood. Scale bar = 100 μm. **e,** Quantification of TUNEL positive cells per midgut at 2 hours post *in vivo* infection (*n* (MOYO-R naive) = 20; *n* (MOYO-R DENV-2) = 21; *n* (MOYO-S naive) = 23; *n* (MOYO-S DENV-2) = 24). The horizontal line indicates the median. Treatments without a common letter were found to be statistically significant (α = 0.05) as calculated by Kruskal-Wallis test with Mann-Whitney post hoc comparison (Kruskal-Wallis chi-squared = 41.907, df = 3, p-value = 4.199e-09).

We hoped to use Z-Vad-FMK or another small molecule caspase inhibitor to confirm the role of caspase activation in the RIA phenotype observed by TUNEL. However, addition of DMSO solvent alone in the *ex vivo* infection media reduced the appearance of TUNEL-positive cells after virus infection (**Extended Data Figure 2)**.

To see the effect of a lack of RIA on the course of infection, we used human Alpha-1 anti-trypsin (hAAT) as a water-soluble apoptosis inhibitor. hAAT is a serum protein serine protease, which has also been found to inhibit caspase-3 activation and subsequently suppress apoptosis in human cells^31^. hAAT is also involved in the human acute infection response as up to a 4-fold increase in hAAT serum concentration was found following stimulation of innate immune cells in donor blood with heat killed *Staphylococcus epidermidis*^32^, although induction of hAAT has not been observed in human flavivirus infection^33^.

We found that supplementation of the infective blood meal with 10 mg/mL of clinical grade hAAT (serum level of hAAT in healthy subjects is around 1.5-3.5 mg/mL) suppressed RIA following exposure to ZIKV or DENV-2, as evidenced by a significant decrease in the number of TUNEL positive cells in hAAT-supplemented mosquitoes (p = 4.62e-06, and 2.38e-05, respectively) (**Fig. 4b,c**). The suppression of apoptosis by hAAT supplementation is short-lived, with levels of TUNEL positive cells in supplemented mosquitoes returning to control infection levels by 24 hpi (**Extended Data Figure 3).** This implies that the effect of apoptosis inhibition is specifically due to RIA at early infection timepoints.

**Fig. 4:**
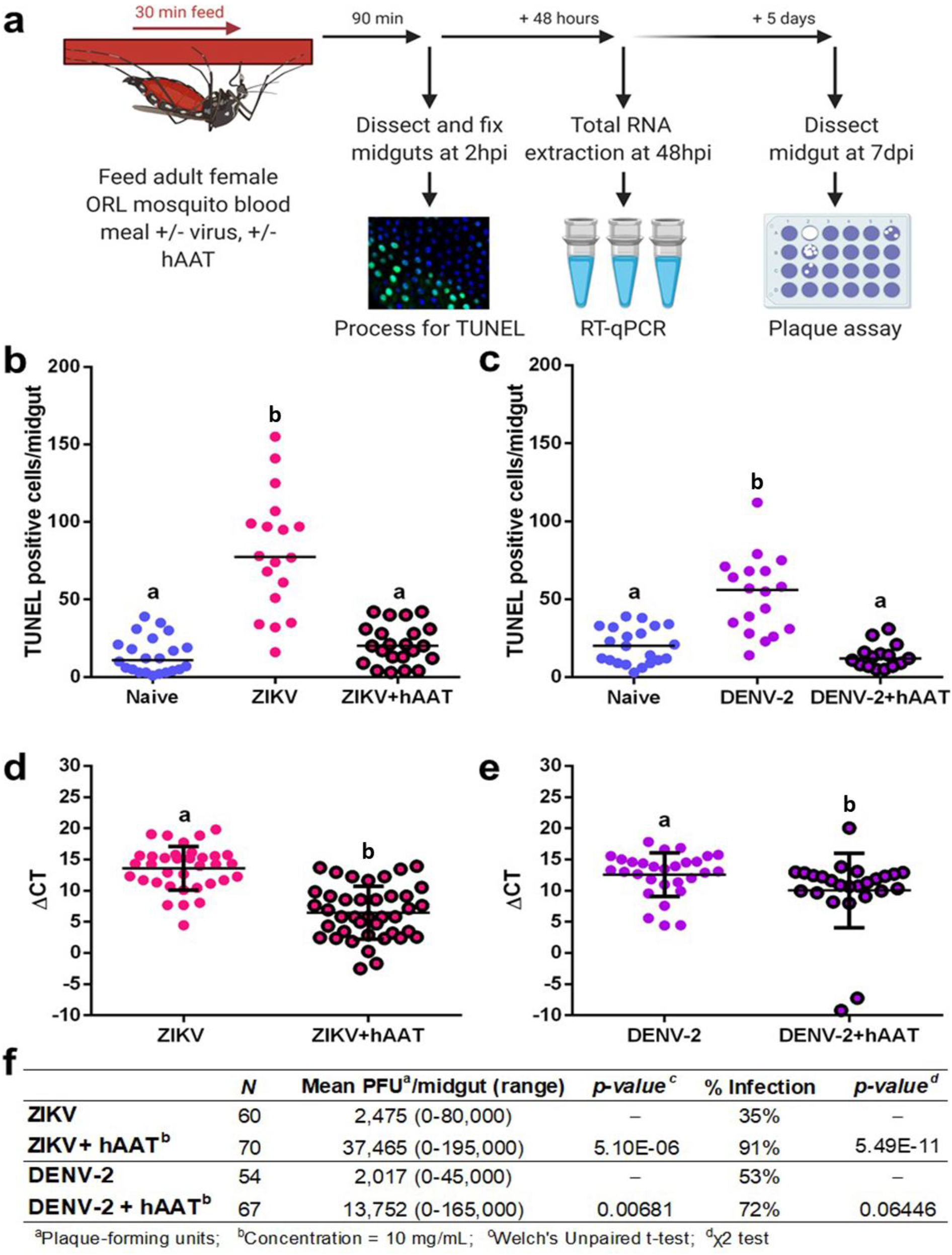
Supplementing a DENV-2 or ZIKV bloodmeal with hAAT in adult female *Aedes aegypti* (ORL) mosquitoes inhibits the rapid induction of apoptosis in the midgut at 2hpi and increases subsequent midgut viral replication. **a,** Workflow of the apoptosis inhibitor co-feeding experiment, hpi=hours post infection, dpi=days post infection. **b-c,** TUNEL positive cell counts in midguts from mosquitoes fed a **(b)** ZIKV (*n* (naive) = 22; *n* (ZIKV) = 18; *n* (ZIKV + hAAT) = 22) or **(c)** DENV-2 infected blood meal or a virus infected blood meal supplemented with 10 mg/mL hAAT (*n* (naive) = 21; *n* (DENV-2) = 18; *n* (DENV-2 + hAAT) = 15) . Treatments without a common letter were found to be statistically significant (α = 0.05) as calculated by Kruskal-Wallis test with Mann-Whitney post-hoc test (panel a: Kruskal-Wallis chi-squared = 32.532, df = 2, p-value = 8.624e-08; panel b: Kruskal-Wallis chi-squared = 26.513, df = 2, p-value = 1.749e-06) **d-e,** ΔCT values from RT-qPCR detecting **(d)** ZIKV (*n* (ZIKV) = 34; *n* (ZIKV + hAAT) = 40) or **(e)** DENV-2 genome copy in whole mosquitoes 48 hours after an infected blood meal with or without 10 mg/mL hAAT (*n* (DENV-2) = 29; *n* (DENV-2 + hAAT) = 25). The horizontal line indicates the median. Treatments without a common letter were found to be statistically significant (α = 0.05) as calculated by Mann-Whitney U test (panel d W = 905, p-value = 1.735e-08; panel e W = 466, p-value = 0.006808). **f,** Productive midgut infection quantified by midgut plaque assay at 7 days after feeding on a ZIKV or DENV-2 infected blood meal (*n* (ZIKV) = 34; *n* (ZIKV + hAAT) = 40; *n* (DENV-2) = 29; *n* (DENV-2 + hAAT) = 25)). Treatments without a common letter were found to be statistically significant (α = 0.05) as calculated by Mann-Whitney U test (ZIKV W = 608, p-value = 1.102e-12; DENV-2 W = 1228.5, p-value = 0.001929) .

Addition of hAAT also increased viral replication in this infection model. hAAT supplementation caused a significant reduction in mean difference in cycle threshold (ΔCT) from 13.59 to 6.47 in ZIKV infected mosquitoes (p = 1.74e-08). This reduction was less pronounced but still significant (12.59 to 10.03) in DENV-2 infected mosquitoes (p = 0.00681) (**Fig. 4d,e**). At 7 days post-infection (dpi), significantly more mosquitoes in the hAAT-supplemented ZIKV group were virus positive in the midgut (91% vs 35% for ZIKV, p = 5.49e-11) (**Fig. 4f**). However, there was no significant difference in the number of virus positive mosquitoes in the DENV-2 hAAT-supplemented group (72% vs 54%, p = 0.0645) (**Fig. 4f**). Furthermore, infected hAAT-supplemented mosquitoes had higher titers of infectious virus in the midgut than non-supplemented mosquitoes, with a 15.1-fold increase in mean PFU/midgut in ZIKV infected mosquitoes (p = 1.10e-12) and a 6.8-fold increase in DENV-2 infected mosquitoes (p = 0.00193) (**Fig. 4f**). Since susceptibility to infection could be increased by potential adverse side effects of hAAT supplementation, we also recorded mosquito mortality at the 7 dpi harvest timepoint. There was no observable correlation between mortality and hAAT supplementation (for ZIKV p = 0.183, for DENV-2 p = 0.637, by Chi-squared test) (**Extended Data Table 1**).

Whether the correlation between a virus-refractory phenotype and a strong RIA response holds true in a broader panel of strains is an interesting topic for future investigation. Several studies have observed increased expression of pro-apoptotic genes correlated with refractory phenotypes in *Ae. aegypti* strains^18,20,34^. A previous study reported induction of the pro-apoptotic gene *Michelob_x* (*mx*), detected by qRTPCR, at 3 hpi with DENV-2 (Jamaica 1409 strain) in the midgut of MOYO-R but not MOYO-S^27^.

We selected mx, IMP, and DNR1 as upstream regulators of caspase activation which have previously been implicated in the innate immune response and may be involved in triggering RIA. *Mx* and *IMP* are the characterized members of the inhibitor of apoptosis (IAP)-antagonist gene family in *Aedes aegypti*, which induce apoptosis by alleviating inhibition of caspases by IAPs^35^. In our ORL infection model, transcript level of *mx* was significantly increased in ZIKV fed mosquitoes but not in DENV-2 fed mosquitoes. *IMP* transcript level was not impacted by infection status **(Extended Data Figure 4)**.

Bioinformatic analyses^36^ of the recently updated *Ae. aegypti* genome (version AaegL5)^37^ suggested that additional IAP-antagonists are present. The presence of these multiple IAP-antagonists could be imperative for the RIA response, as deleting the regulatory region responsible for coordinated induction of three IAP-antagonist genes (*reaper*, *sickle*, and *hid*) in *Drosophila* adults significantly suppressed RIA and rendered the flies more susceptible to FHV^27^. Whether an orthologous regulatory region in *Aedes aegypti* is present and responsible for mediating the RIA response to the flaviviruses tested herein remains to be seen. The differential RIA response observed in *Ae. aegypti* strains could be due to genetic polymorphisms present in enhancer or regulatory regions as polymorphisms in cis-regulatory elements between MOYO-R and MOYO-S have previously been observed^38^.

Defense repressor 1 (DNR1) is a negative apoptosis regulator, which has been characterized in *Aedes aegypti, Aedes albopictus* cells and *Drosophila* as depleting levels of the caspase dronc by ubiquitination^18,39,40^. DNR1 is also a negative regulator of the immune deficiency (IMD) pathway in *Drosophila*^41^ and is depleted following Sindbis virus infection of *Aedes albopictus* cells^40^, making it a gene of interest for investigating cross-talk between innate immunity and apoptosis. In this infection model, there was no change in transcript level of *DNR1* at the 2hpi timepoint between naive blood and virus-fed mosquitoes **(Extended Data Figure 4)**. In the future, studies investigating protein-level regulation of these genes during the RIA response should be undertaken to confirm involvement or lack thereof.

Studies in *Drosophila* indicated that the transcriptional regulation of key pro-apoptotic genes is influenced by epigenetic modification of the regulatory block harboring the IAP-antagonist genes ^42,43^. The epigenetic landscape of this region is controlled by *cis* DNA elements^44^ and is responsive to environmental cues^45^. If a similar mechanism is conserved in mosquitoes, the differential RIA response observed in *Ae. aegypti* strains could also be due to genetic polymorphisms present in regulatory regions controlling the apoptotic epigenetic landscape.

An interesting feature of the RIA in our infection models was the clustering of TUNEL-positive cells. There is no previous evidence suggesting a clustered sub-population of cells in the midgut epithelium would be particularly susceptible to DENV-2 or ZIKV infection. Therefore, while the presence of virus in any or all cells undergoing RIA (i.e., TUNEL-positive) remains to be seen, we speculate that cells adjacent to those infected may also undergo apoptosis **(Fig. 5)**. In mammalian systems, virus infection-induced apoptosis of uninfected cells has been observed in Herpes Simplex Virus-1 and Human Immunodeficiency virus infection^46,47^. In plants, the rapid induction of cell death following pathogen infection often involves cells surrounding the infected cells ^48,49^. This type of cell death, termed Hypersensitive Response (HR), is responsible for mediating resistance to pathogen infection in higher plants. The mechanism of HR in plants has not been completely understood, but likely involves the production and release of reactive oxygen species^50,51^. Apoptosis-induced-apoptosis, in which a cell undergoing apoptosis stimulates apoptosis in its neighbors by paracrine signaling, has been observed in *Drosophila* and is mediated by Eiger, the insect ortholog of mammalian tumor necrosis factor^52^. An Eiger ortholog (AAEL010524-PA) has been identified in *Ae. aegypti*. Previous research on West Nile Virus infected *Culex quinquefasciatus* cells suggests that paracrine signaling from infected cells induces an antiviral state in their neighbors via activation of the JAK-STAT pathway^53^. The presence of pro-apoptotic signaling and the relationship of RIA to known immune pathways in *Ae. aegypti* is an attractive topic of future study.

**Fig. 5:**
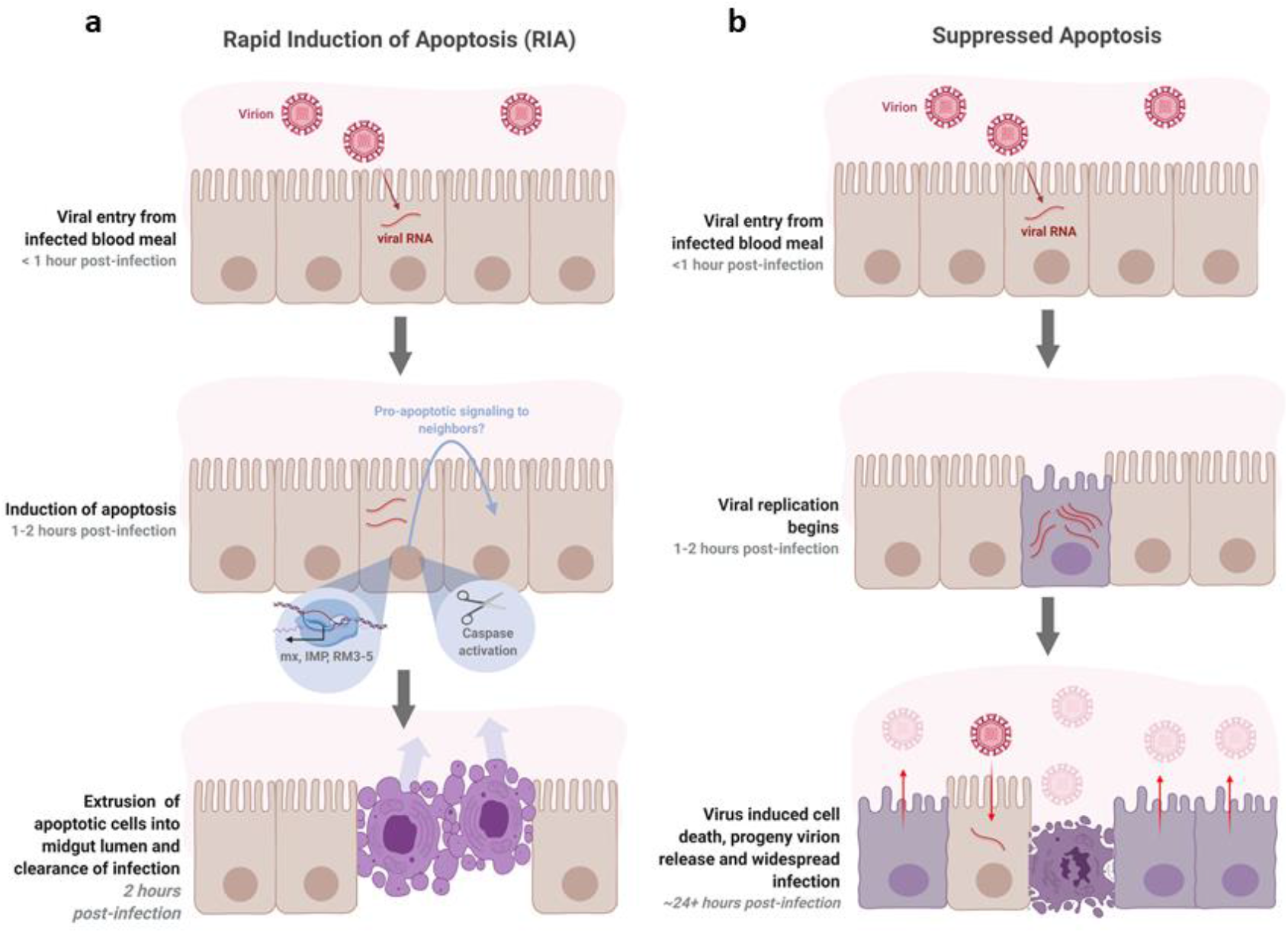
Hypothesized model of antiviral apoptosis in the *Aedes aegypti* midgut. **a,** RIA of infected cells in the midgut epithelium within 2 hours of an infected blood meal arrests viral replication before progeny virions can be produced and suppresses infection. The clustered appearance of apoptotic cells may be caused by pro-apoptotic signaling to neighboring cells. **b,** At later timepoints post-infection, when infection is widespread throughout the midgut and progeny virions are being produced, death of infected cells may facilitate release of virions from cells and/or from the midgut by degrading the integrity of the epithelium and basal lamina.

In summary, we found that *Ae. aegypti* midgut epithelial cells undergo RIA within 2 hours of exposure to DENV-2 and ZIKV, and that this response is likely caspase mediated. We noted that inhibiting RIA via hAAT treatment increased virus proliferation in the midgut. We propose that RIA beginning 1-4 hpi is a mosquito response to virus exposure distinct from 247 blood meal- or virus-induced cell death that occurs at later times post-infection. The increase in infection severity upon suppression of RIA suggests this phenomenon plays a significant role in mediating the midgut infection barrier in vector mosquitoes **(Fig. 5)**.

## Supporting information

Raw Source Data

## Acknowledgements

The authors thank Dr. Sihong Song for discussion on hAAT and providing us clinical grade hAAT and the United States Department of Agriculture’s Center for Medical, Agricultural and Veterinary Entomology for the *Aedes aegypti* Orlando strain. The authors also thank Dr. David W. Severson and his group for hypothesis generation and collaboration on previous work in this vein as well as for sharing the MOYO strains, and Alyssa Cornista and Ashley Gomez for technical support. This research was supported in part by the United States Centers for Disease Control (CDC) Grant 1U01CK000510-03: Southeastern Regional Center of Excellence in Vector-Borne Diseases: The Gateway Program. The CDC had no role in the design of the study, the collection, analysis, and interpretation of data, or in writing the manuscript. Support was also provided by the University of Florida Emerging Pathogens Institute and the University of Florida Preeminence Initiative (RRD) and NIH GM106174 (LZ). The following reagents were obtained through BEI Resources, NIAID, NIH: Purified Zika Virus, PRVABC59, NR-50684 and Dengue Virus Type 2, New Guinea C, NR-51427. Workflow figures and summary figure were created with Biorender.com.

## MATERIALS AND METHODS

### Statistical analysis

Statistical analyses were performed using RStudio Desktop, Open Source Edition and R version 4.0.0^54^. All experiments analyzed represent pooled data from 3 independent replicates. Data was tested for normality using Shapiro-Wilk tests. For TUNEL positive cell count experiments, statistical significance was analyzed via Kruskal-Wallis omnibus test with Mann-Whitney pairwise post hoc test. Infection intensity data from plaque assay and RT-qPCR results were analyzed by Mann-Whitney test. Differences in infection prevalence from plaque assay data were analyzed by a chi-squared test. All hypothesis testing was performed at α=0.05.

### Virus culture

Vero C1008 (clone E6, ATCC) African green monkey kidney cells were grown in complete Dulbecco’s Modified Eagle Medium (10% FBS) until 90% confluent, then transferred to reduced DMEM (3% FBS) for 24 hours before inoculation. Virus was allowed to adsorb for 2 hours at 37°C, after which media was replaced with non-infectious reduced DMEM. Flasks were observed every 3 days. The supernatant was harvested when 50% of the cells showed visible signs of flavivirus induced cytopathic effects (CPE). Cell debris was spun out of the stock, which was then stabilized with 15% trehalose and stored in liquid nitrogen. Titration of stock virus was done by plaque assay on baby hamster kidney cells (BHK-21 Clone 13, ATCC). Strains used in this study were: DENV-2 New Guinea C strain (passage 3, 10^6^ PFU/mL; BEI resources) and ZIKV Puerto Rico PRVABC59 (passage 3, 10^6^ PFU/mL; BEI resources).

### Mosquito rearing

All mosquito strains (MOYO-R, MOYO-S, and Orlando) were housed in 27°C growth chambers at 80% humidity on a 12/12 light/dark cycle. Eggs were hatched in warmed deionized (DI) water. Larvae were grown at a density of 400 per 18” × 12.5” larval tray and were fed a 50:50 mix of beef liver powder and ground TetraMin^®^ fish flakes. Pupae were removed daily using plastic transfer pipets. Adults were maintained on a 10% sucrose solution.

### *In vivo* infection

Live mosquito infection was performed under Biosafety Level (BSL) 2+ practice in an Arthropod Containment Level 3 facility. Virus-infected blood meals were a 2:2:1 mixture of O+ human RBCs, 10^6^ PFU/mL virus stock, and heat inactivated human serum. For naive blood meals, the virus stock was replaced by Vero E6 culture supernatant with 15% trehalose. Mosquitoes were fasted on DI water for 12-24 hours prior to blood-feeding and blood fed using a water-jacketed membrane feeder maintained at 37°C for 30 minutes. For experiments including MOYO-R or MOYO-S mosquitoes, pork sausage casing was used as the membrane, while parafilm was used for ORL mosquitoes. After blood feeding, mosquitoes were cold anesthetized (−20°C) and those that had not blood fed were killed and discarded. At the harvest timepoint, mosquitoes were cold anaesthetized, rinsed 1x in 50% EtOH, and 2x in PBS, and dissected in PBS. Dissected midguts were immediately transferred to 4% paraformaldehyde in PEMS fixation buffer (10 mM EGTA, 1 mM MgSO_4_, 100 mM PIPES (pH 6.9), 75 mM sucrose, 0.1% Triton X-100, in H_2_O).

### hAAT treatment

Clinical grade hAAT (Grifol’s therapeutics, Catalog # NDC 13533-702-11) was dissolved in autoclaved DI water and added to the blood meal or inoculum at a concentration of 10 mg/mL. The volume of drug was subtracted from the volume of serum added. Negative controls had an equivalent volume of serum or stock replaced by autoclaved DI water.

### *Ex vivo* infection

*Ex vivo* infections were performed in a BSL-2 facility. Non-bloodfed female mosquitoes were dissected in plain DMEM and 10 midguts per infection were pooled, then transferred to virus stock (or Vero E6 cell culture supernatant with 15% trehalose for negative controls) diluted 2:3 into plain DMEM to match the concentration of virus in blood meals for the *in vivo* infection. Midguts were incubated at room temperature on a rocker for 1-2 hours as the experiment required.

### Midgut plaque assay

Midguts were dissected, submerged in 150 μL reduced serum DMEM (3% FBS) and stored at −80°C. After thawing, midguts were homogenized using a Next Advance Bullet Blender 24, and debris was pelleted. Supernatant was serially diluted 10-fold out to 10^3^ in reduced serum DMEM and applied to baby hamster kidney cells (BHK-21 Clone 13, ATCC) for 1 hour. Inoculum was replaced with 0.8% methylcellulose in reduced serum DMEM and cells were incubated for 4 days before crystal violet stain was applied. Plaques were counted manually.

### Viral genome quantification

RNA was extracted from individual whole mosquitoes stored in 150μL Trizol^®^ reagent (Thermo Fisher Scientific Catalog # 15596018) using a Zymo Quick-RNA™ MiniPrep kit (Catalog # R1055). cDNA was prepared from 1000 ng of RNA using a high capacity cDNA reverse transcription kit (Applied Biosystems Ref# 4368814) and a thermocycling program of 25°C for 10 minutes, 37°C for 2 hours, and 85°C for 5 min, followed by a 4°C indefinite hold. qPCR was performed in duplicate for each target using an Applied Biosystems 7500fast Real-Time PCR System. Primer information can be found in Extended Data Table 2. Primers were designed in house using Primer 3^55^, except *AeDNR1* primers which were sourced from a previous study^18^. Amplicons were sequenced to confirm specificity. The thermocycler program used was 50°C for 20 seconds, 95°C for 10 min; followed by 40 cycles of 95°C for 15 seconds and 60°C for 1 min; followed by a melting curve stage (15 seconds at 95^°^C, 1 min at 60°C, 30 seconds at 95°C, 15 seconds at 60°C). Delta-CT (ΔCT) values were calculated by subtracting the RPL32 CT value from the viral genome CT value. Relative expression values were calculated as 2^−ΔΔCT^, where ΔΔCT was found by subtracting the arithmetic mean ΔCT of the naive blood control across all replicates from each sample ΔCT.

### Terminal deoxynucleotidyl transferase dUTP nick end labeling (TUNEL)

Pools of 7-10 midguts were fixed in 4% paraformaldehyde in 5 mM PIPES, pH 7.2 5 mM NaCl, 5 mM MgCl2, 1 mM EGTA (PME buffer) for 30 minutes. They were dehydrated in 25%-50%-75%-100% methanol in a Tris buffered saline (TBS) gradient with 5 minutes in each step and stored at 4°C in 100% methanol until processing. Midguts were rehydrated in the reverse gradient. If midguts were from recently blood-fed mosquitoes, they were split open and flattened with a tungsten needle so the blood bolus could be removed. Midguts were then permeabilized in 20 μg/mL proteinase K for 5 minutes at room temperature, then rinsed 2x, washed for 5 minutes 2x, and fixed for 10 minutes in 4% paraformaldehyde in TBS. TUNEL reactions were performed using equilibration buffer and labelling mix from the Millipore FragEL™ DNA Fragmentation Detection Kit, Fluorescent (FITC conjugated) (QIA39-1EA) used as recommended by the manufacturer with15 U Calf Thymus TdT (Millipore PF060). Midguts were whole mounted for observation on a Zeiss Axioplan 2 imaging microscope, and representative images were taken using a Keyence BZ-X700. TUNEL positive cells were manually counted.

### Pan-caspase activation assay

At 1-hour post infection, inoculum was removed from *ex-vivo* infected guts and replaced with 300 μL of plain DMEM. Red-VAD-FMK (MBL catalog # JM-K190) was added to the media at the concentration recommended by the manufacturer and tissues were incubated for an additional hour at room temperature. Tissues were washed in the provided wash buffer and resuspended in wash buffer for whole mount fluorescence microscopy. A Zeiss Axioplan 2 imaging microscope was used to visualize the assay, and imaging was completed within 20 minutes of the end of the incubation period.

### Data Availability Statement

The authors confirm that the data supporting the findings of this study are available within the supplementary materials of this article.

---

**Extended Data Figure 1:**
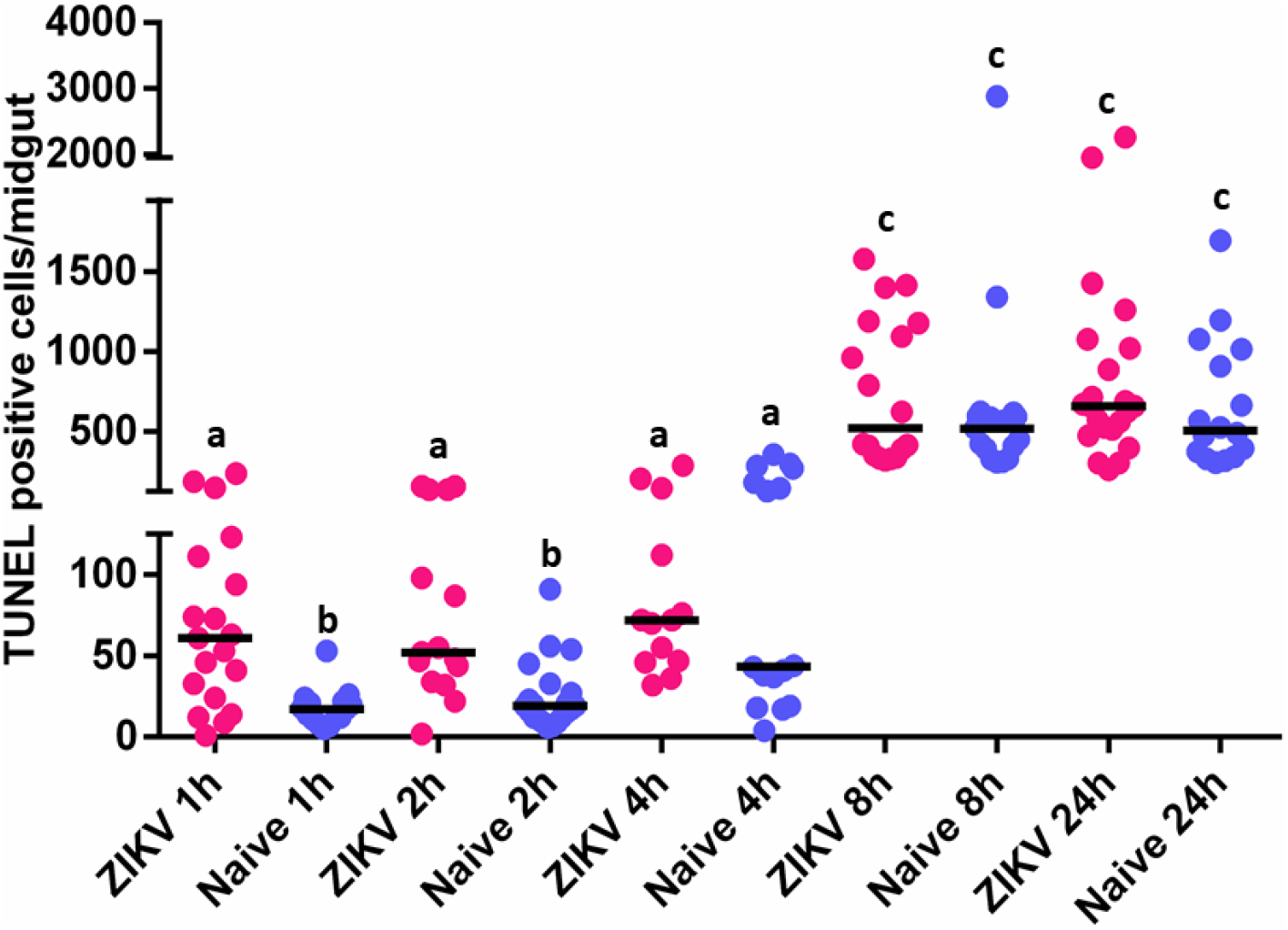
Adult female ORL strain mosquitoes were fed a naive or ZIKV infected bloodmeal. Midguts were dissected and processed for TUNEL at the indicated times post infection. By 8 hours post-feeding, both naive and virus-fed groups have equivalently high levels of apoptotic cells, likely representing normal turnover of the gut epithelium in the process of blood digestion. An increase in TUNEL positive cells in virus fed mosquitoes is seen between 1 and 4 hours post-feeding (*n* (ZIKV 1h) = 19; *n* (naive 1h) = 15; *n* (ZIKV 2h) = 15; *n* (naive 2h) = 21; *n* (ZIKV 4h) = 13; *n* (naive 4h) = 16; *n* (ZIKV 8h) = 18; *n* (naive 8h) = 17; *n* (ZIKV 24h) = 22; *n* (naive 24h) = 15). Data is pooled from 3 independent replicates. Treatments without a common letter were found to be statistically significant (α = 0.05) as calculated by Kruskal-Wallis test with Mann-Whitney post-hoc test.

**Extended Data Table 1:** Adult female ORL mosquitoes were fed a ZIKV or DENV-2 infected blood meal with or without 10 mg/mL hAAT and mortality was assessed at 7 days post infection during harvest for plaque assay. hAAT treatment did not cause any observable increase in mortality. Data is pooled from three replicates and significance was assessed at α=0.05 via a chi-squared test.

|  | *N* | % Mortality | *p*-value<sup>a</sup> |
| --- | --- | --- | --- |
| ZIKV | 78 | 8.5 | – |
| ZIKV+ hAAT<sup>b</sup> | 77 | 12.9 | *p* = 0.1827 |
| DENV-2 | 59 | 15.4 | – |
| DENV-2 + hAAT<sup>b</sup> | 77 | 6.5 | *p* = 0.6368 |
| <sup>b</sup>Concentration=10 mg/mL |  |  |  |
| <sup>a</sup>Calculated by $\chi^2$ | | | |

**Extended Data Figure 2:**
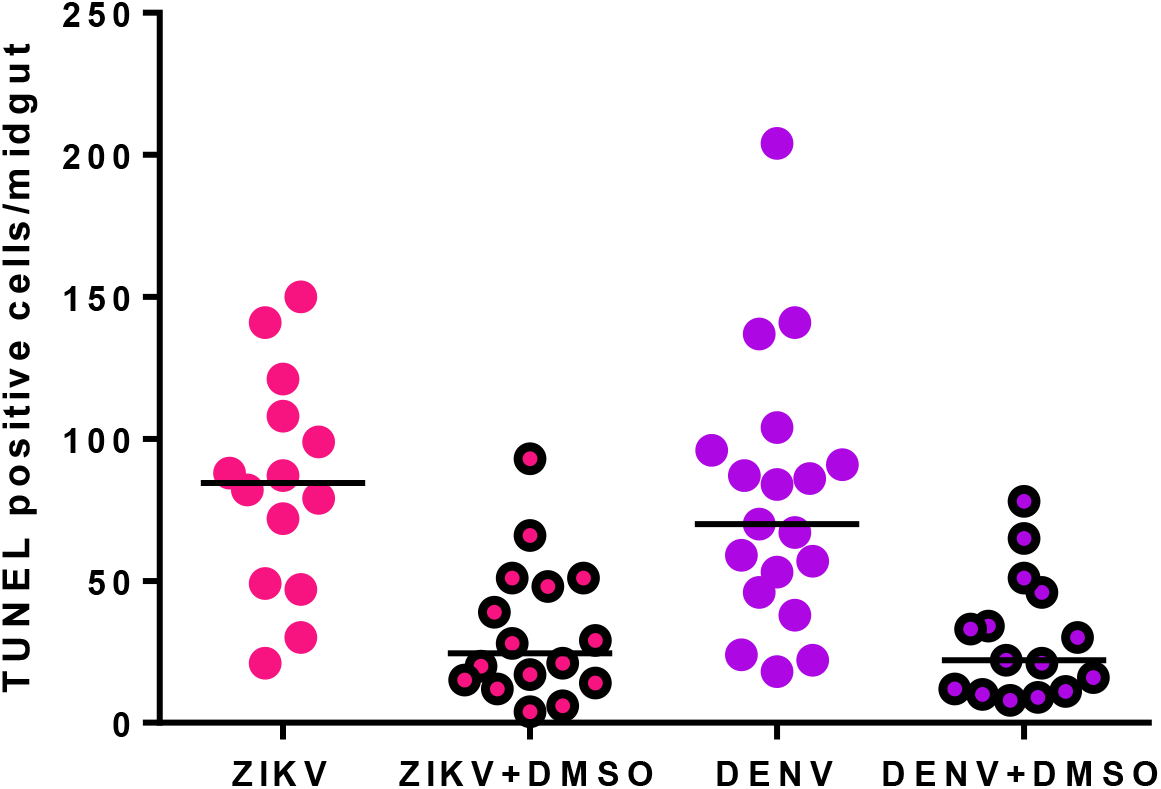
Midguts treated with DMSO vehicle during *ex vivo* infection show reduced rapid induction of apoptosis. Midguts were dissected from adult female non-bloodfed ORL mosquitoes and infected *ex vivo* with plain virus stock or virus stock plus 1% molecular biology grade DMSO (Millipore-Sigma D2438-10mL). Data is pooled from 2 independent replicates. Horizontal line represents median. (*n* (ZIKV) = 14, *n* (ZIKV + DMSO) = 16, *n* (DENV) = 19, *n* (DENV + DMSO) = 15).

**Extended Data Figure 3:**
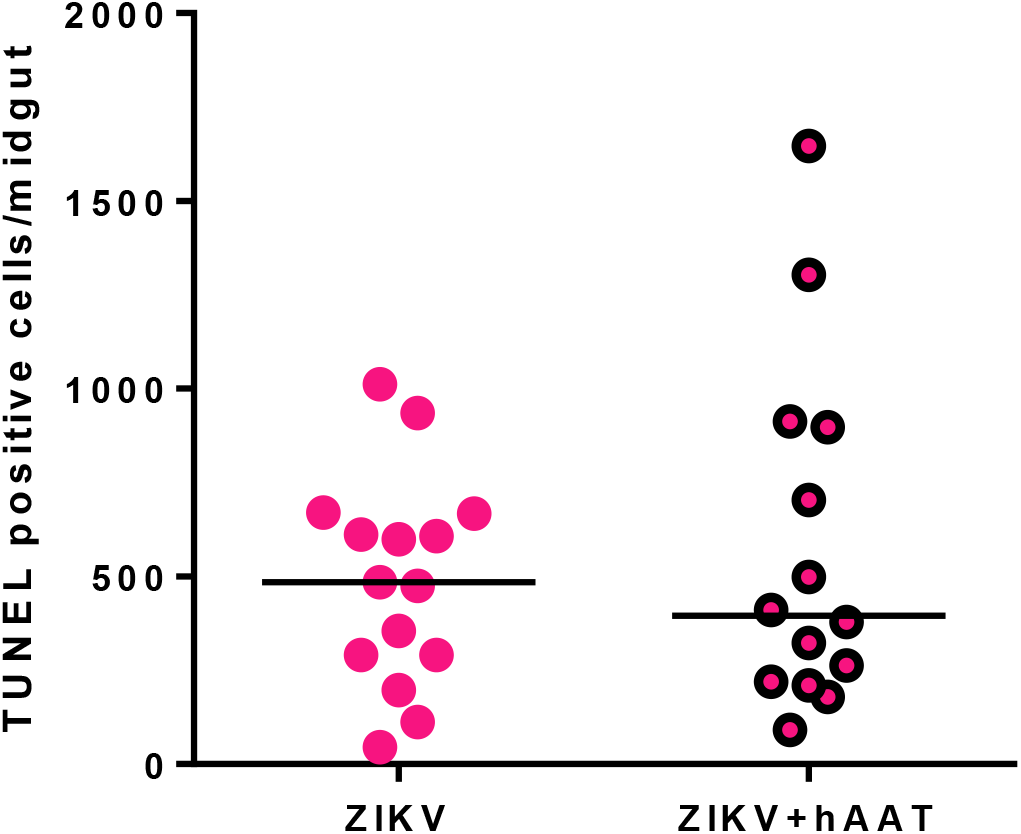
Apoptosis inhibition from hAAT subsides by 24h post-infection. Adult female ORL mosquitoes were fed a blood meal containing 10^6^ PFU/mL with or without 10mg/mL hAAT supplement. Midguts were dissected and prepared for TUNEL at 24h post-infection. Data is pooled from 2 independent replicates. (*n* (naïve) = 15, *n* (ZIKV + hAAT) = 14). Horizontal line indicates median.

**Extended Data Figure 4:**
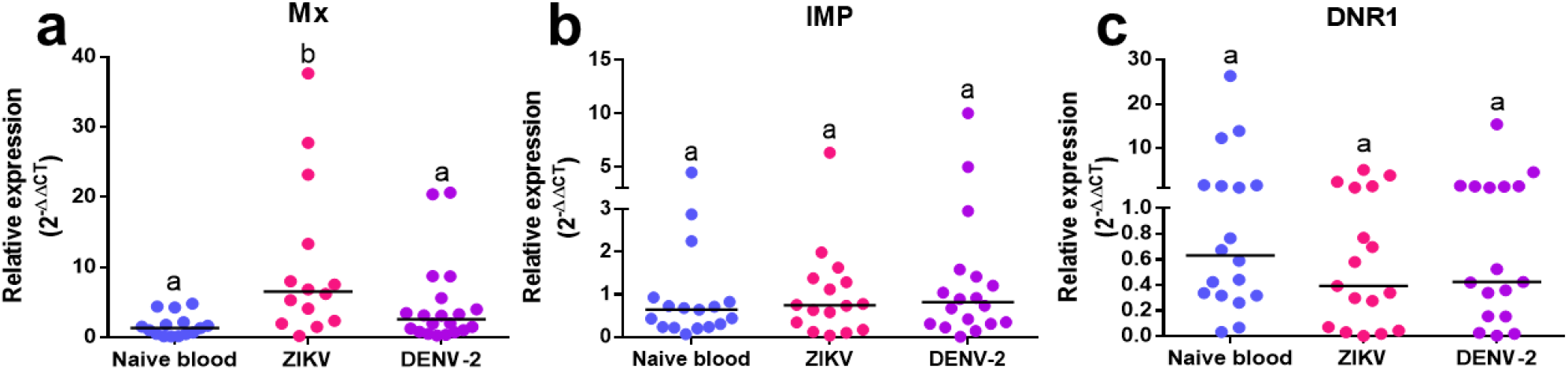
Transcript level of IAP-antagonist *mx* is significantly increased during the RIA response to ZIKV infection, *IMP* and negative apoptosis regulator *DNR1* are not. Pools of 3 midguts were dissected from naive blood, ZIKV, or DENV-2 fed ORL strain female mosquitoes at 2hpi. Transcript expression level of the indicated gene was analyzed by RT-qPCR relative to *RPL32*. Data is pooled from 3 independent replicates (4 replicates in the case of *mx*). Treatments without a common letter were found to be statistically significant (α = 0.05) as calculated by Kruskal-Wallis test with Mann-Whitney post-hoc test. Black line indicates median. (*n* (*mx* naive blood) = 15, *n* (*mx* ZIKV) = 14, *n* (*mx* DENV-2) = 20, *n* (*IMP* naive blood) = 17, *n* (*IMP* ZIKV) = 16, *n* (*IMP* DENV-2) = 18, *n* (*DNR1* naive blood) = 18, *n* (*DNR1* ZIKV) = 17, *n* (*DNR1* DENV-2) = 17).

**Extended Data Table S2:**
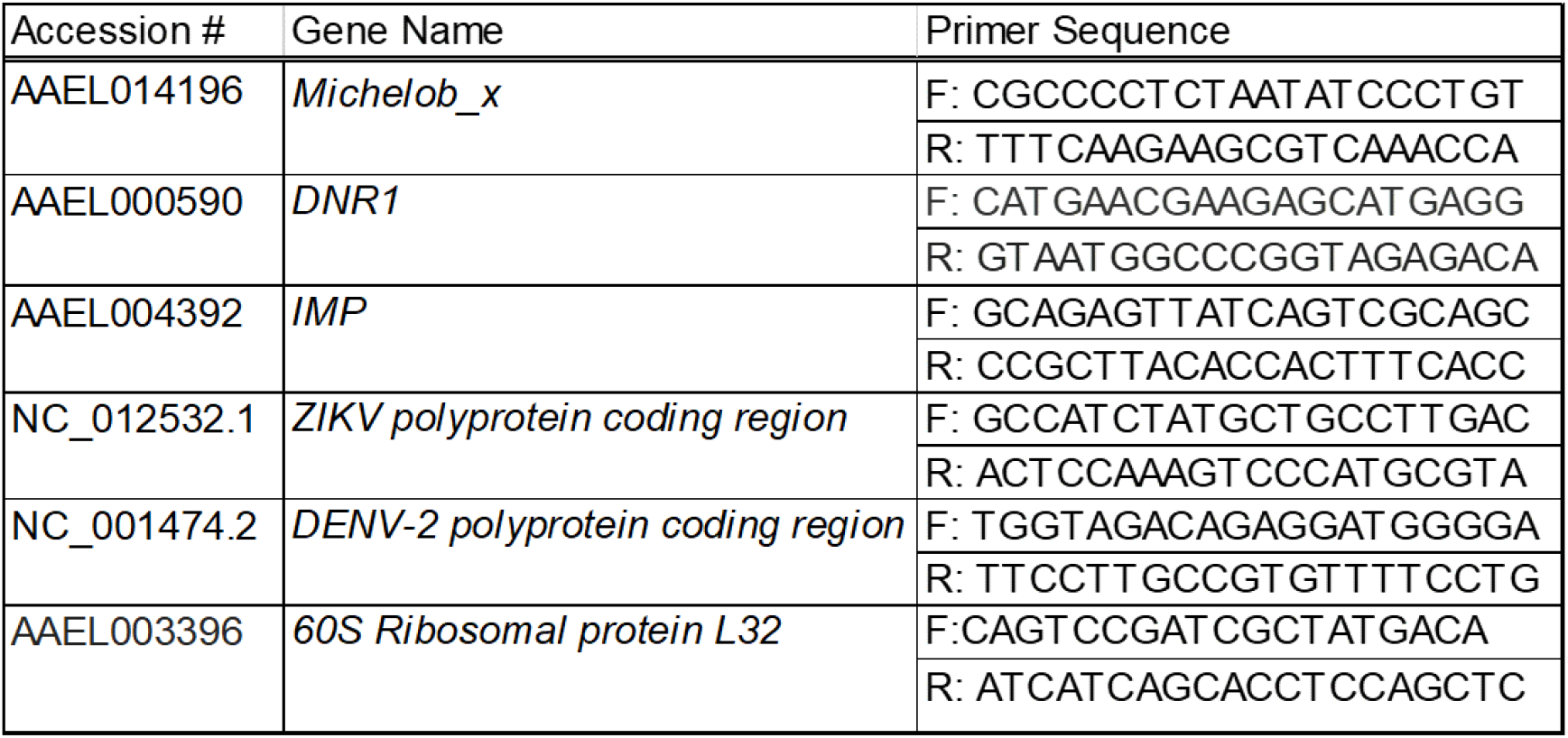
RT-qPCR primer sequences, gene names, and accession numbers.

## Notes

### Competing Interest Statement

The authors have declared no competing interest.

## References

1. Franz, A. W. E., Kantor, A. M., Passarelli, A. L. & Clem, R. J. Tissue barriers to arbovirus infection in mosquitoes. Viruses 7, 3741–3767 (2015).

2. Caicedo, P. A. et al. Selection of *Aedes aegypti* (Diptera: Culicidae) strains that are susceptible or refractory to Dengue-2 virus. Can Entomol 145, 273–282 (2013).

3. Kalayanarooj, S. Clinical manifestations and management of dengue/dhf/dss. Trop. Med. Health 39, 83–87 (2011).

4. Messina, J. P. et al. A global compendium of human dengue virus occurrence. Sci. Data 1, 140004 (2014).

5. Barbi, L., Coelho, A. V. C., Alencar, L. C. A. de & Crovella, S. Prevalence of Guillain-Barré syndrome among Zika virus infected cases: a systematic review and meta-analysis. Braz J Infect Dis 22, 137–141 (2018).

6. Brady, O. J. et al. The association between Zika virus infection and microcephaly in Brazil 2015-2017: An observational analysis of over 4 million births. PLoS Med. 16, e1002755 (2019).

7. Souza-Neto, J. A., Powell, J. R. & Bonizzoni, M. Aedes aegypti vector competence studies: A review. Infect. Genet. Evol. 67, 191–209 (2019).

8. Sim, S. et al. Transcriptomic profiling of diverse Aedes aegypti strains reveals increased basal-level immune activation in dengue virus-refractory populations and identifies novel virus-vector molecular interactions. PLoS Negl. Trop. Dis. 7, e2295 (2013).

9. Angleró-Rodríguez, Y. I. et al. Aedes aegypti Molecular Responses to Zika Virus: Modulation of Infection by the Toll and Jak/Stat Immune Pathways and Virus Host Factors. Front. Microbiol. 8, 2050 (2017).

10. Bennett, K. E. et al. Variation in vector competence for dengue 2 virus among 24 collections of Aedes aegypti from Mexico and the United States. Am. J. Trop. Med. Hyg. 67, 85–92 (2002).

11. Roulston, A., Marcellus, R. C. & Branton, P. E. Viruses and apoptosis. Annu. Rev. Microbiol. 53, 577–628 (1999).

12. Upton, J. W. & Chan, F. K.-M. Staying alive: cell death in antiviral immunity. Mol. Cell 54, 273–280 (2014).

13. Everett, H. & McFadden, G. Apoptosis: an innate immune response to virus infection. Trends Microbiol. 7, 160–165 (1999).

14. Clem, R. J. Viral IAPs, then and now. Semin. Cell Dev. Biol. 39, 72–79 (2015).

15. Clem, R. J., Hardwick, J. M. & Miller, L. K. Anti-apoptotic genes of baculoviruses. Cell Death Differ. 3, 9–16 (1996).

16. Slonchak, A. et al. Zika virus noncoding RNA suppresses apoptosis and is required for virus transmission by mosquitoes. Nat. Commun. 11, 2205 (2020).

17. O’Neill, K., Olson, B. J. S. C., Huang, N., Unis, D. & Clem, R. J. Rapid selection against arbovirus-induced apoptosis during infection of a mosquito vector. Proc. Natl. Acad. Sci. USA 112, E1152–61 (2015).

18. Eng, M. W., van Zuylen, M. N. & Severson, D. W. Apoptosis-related genes control autophagy and influence DENV-2 infection in the mosquito vector, Aedes aegypti. Insect Biochem. Mol. Biol. 76, 70–83 (2016).

19. Girard, Y. A. et al. Salivary gland morphology and virus transmission during long-term cytopathologic West Nile virus infection in *Culex* mosquitoes. Am. J. Trop. Med. Hyg. 76, 118–128 (2007).

20. Ocampo, C. B. et al. Differential expression of apoptosis related genes in selected strains of Aedes aegypti with different susceptibilities to dengue virus. PLoS One 8, e61187 (2013).

21. Girard, Y. A., Popov, V., Wen, J., Han, V. & Higgs, S. Ultrastructural study of West Nile virus pathogenesis in Culex pipiens quinquefasciatus (Diptera: Culicidae). J. Med. Entomol. 42, 429–444 (2005).

22. Wang, H., Gort, T., Boyle, D. L. & Clem, R. J. Effects of manipulating apoptosis on Sindbis virus infection of Aedes aegypti mosquitoes. J. Virol. 86, 6546–6554 (2012).

23. Romoser, W. S. et al. Pathogenesis of Rift Valley fever virus in mosquitoes--tracheal conduits & the basal lamina as an extra-cellular barrier. Arch. Virol. Suppl. 89–100 (2005). doi:10.1007/3-211-29981-5_8

24. Vogels, C. B., Göertz, G. P., Pijlman, G. P. & Koenraadt, C. J. Vector competence of European mosquitoes for West Nile virus. Emerg. Microbes Infect. 6, e96 (2017).

25. Means, J. C. & Passarelli, A. L. Viral fibroblast growth factor, matrix metalloproteases, and caspases are associated with enhancing systemic infection by baculoviruses. Proc. Natl. Acad. Sci. USA 107, 9825–9830 (2010).

26. Bideshi, D. K., Tan, Y., Bigot, Y. & Federici, B. A. A viral caspase contributes to modified apoptosis for virus transmission. Genes Dev. 19, 1416–1421 (2005).

27. Liu, B. et al. P53-mediated rapid induction of apoptosis conveys resistance to viral infection in Drosophila melanogaster. PLoS Pathog. 9, e1003137 (2013).

28. Okuda, K. et al. Cell death and regeneration in the midgut of the mosquito, Culex quinquefasciatus. J. Insect Physiol. 53, 1307–1315 (2007).

29. Yan, G., Christensen, B. M. & Severson, D. W. Comparisons of genetic variability and genome structure among mosquito strains selected for refractoriness to a malaria parasite. J. Hered. 88, 187–194 (1997).

30. Schneider, J. R., Mori, A., Romero-Severson, J., Chadee, D. D. & Severson, D. W. Investigations of dengue-2 susceptibility and body size among Aedes aegypti populations. Med Vet Entomol 21, 370–376 (2007).

31. Zhang, B. et al. Alpha1-antitrypsin protects beta-cells from apoptosis. Diabetes 56, 1316–1323 (2007).

32. Pott, G. B., Chan, E. D., Dinarello, C. A. & Shapiro, L. Alpha-1-antitrypsin is an endogenous inhibitor of proinflammatory cytokine production in whole blood. J. Leukoc. Biol. 85, 886–895 (2009).

33. Kunder, M., Lakshmaiah, V. & Moideen Kutty, A. V. Plasma Neutrophil Elastase, α1-Antitrypsin, α2-Macroglobulin and Neutrophil Elastase-α1-Antitrypsin Complex Levels in patients with Dengue Fever. Indian J. Clin. Biochem. 33, 218–221 (2018).

34. Serrato, I. M., Caicedo, P. A., Orobio, Y., Lowenberger, C. & Ocampo, C. B. Vector competence and innate immune responses to dengue virus infection in selected laboratory and field-collected Stegomyia aegypti (= Aedes aegypti). Med Vet Entomol 31, 312–319 (2017).

35. Wang, H. & Clem, R. J. The role of IAP antagonist proteins in the core apoptosis pathway of the mosquito disease vector Aedes aegypti. Apoptosis 16, 235–248 (2011).

36. Zhou, L. et al. Michelob_x is the missing inhibitor of apoptosis protein antagonist in mosquito genomes. EMBO Rep. 6, 769–774 (2005).

37. Matthews, B. J. et al. Improved reference genome of *Aedes aegypti* informs arbovirus vector control. Nature 563, 501–507 (2018).

38. Behura, S. K. et al. High-throughput cis-regulatory element discovery in the vector mosquito Aedes aegypti. BMC Genomics 17, 341 (2016).

39. Primrose, D. A. et al. Interactions of DNR1 with the apoptotic machinery of Drosophila melanogaster. J. Cell Sci. 120, 1189–1199 (2007).

40. Li, X. et al. Identification of Aadnr1, a novel gene related to innate immunity and apoptosis in Aedes albopictus. Gene 587, 18–26 (2016).

41. Guntermann, S., Primrose, D. A. & Foley, E. Dnr1-dependent regulation of the Drosophila immune deficiency signaling pathway. Dev. Comp. Immunol. 33, 127–134 (2009).

42. Zhang, Y. et al. Epigenetic blocking of an enhancer region controls irradiation-induced proapoptotic gene expression in Drosophila embryos. Dev. Cell 14, 481–493 (2008).

43. Arya, R. et al. A Cut/cohesin axis alters the chromatin landscape to facilitate neuroblast death. Development 146, (2019).

44. Lin, N. et al. A barrier-only boundary element delimits the formation of facultative heterochromatin in Drosophila melanogaster and vertebrates. Mol. Cell. Biol. 31, 2729–2741 (2011).

45. Zhang, C. et al. An intergenic regulatory region mediates Drosophila Myc-induced apoptosis and blocks tissue hyperplasia. Oncogene 34, 2385–2397 (2015).

46. Stanziale, S. F. et al. Infection with oncolytic herpes simplex virus-1 induces apoptosis in neighboring human cancer cells: a potential target to increase anticancer activity. Clin. Cancer Res. 10, 3225–3232 (2004).

47. Meyaard, L. et al. Programmed death of T cells in HIV-1 infection. Science 257, 217–219 (1992).

48. Ryerson, D. E. & Heath, M. C. Cleavage of Nuclear DNA into Oligonucleosomal Fragments during Cell Death Induced by Fungal Infection or by Abiotic Treatments. Plant Cell 8, 393–402 (1996).

49. Del Pozo, O. & Lam, E. Caspases and programmed cell death in the hypersensitive response of plants to pathogens. Curr. Biol. 8, 1129–1132 (1998).

50. Mur, L. A. J., Kenton, P., Lloyd, A. J., Ougham, H. & Prats, E. The hypersensitive response; the centenary is upon us but how much do we know? J. Exp. Bot. 59, 501–520 (2008).

51. Balint-Kurti, P. The plant hypersensitive response: concepts, control and consequences. Mol. Plant Pathol. 20, 1163–1178 (2019).

52. Pérez-Garijo, A., Fuchs, Y. & Steller, H. Apoptotic cells can induce non-autonomous apoptosis through the TNF pathway. Elife 2, e01004 (2013).

53. Paradkar, P. N., Trinidad, L., Voysey, R., Duchemin, J.-B. & Walker, P. J. Secreted Vago restricts West Nile virus infection in Culex mosquito cells by activating the Jak-STAT pathway. Proc. Natl. Acad. Sci. USA 109, 18915–18920 (2012).

54. R Core Team. R: A language and environment for statistical computing. (R Foundation for Statistical Computing, 2020).

55. Koressaar, T. & Remm, M. Enhancements and modifications of primer design program Primer3. Bioinformatics 23, 1289–1291 (2007).

